# A comparison among EL-FAME, PLFA, and quantitative PCR methods to detect changes in the abundance of soil bacteria and fungi

**DOI:** 10.1101/2024.05.09.593327

**Authors:** José A. Siles, Roberto Gómez-Pérez, Alfonso Vera, Carlos García, Felipe Bastida

## Abstract

EL-FAME (ester-linked fatty acid methyl ester), PLFA (phospholipid fatty acid) and qPCR (quantitative PCR) of ribosomal genes are three of the most common methods used to quantify soil microbial communities. The reliability of these three methods has not been simultaneously compared in situations of rapid changes in soil microbial abundances. For this purpose, we (i) incubated badland, cropland, and forest soils with nutrients or antibiotics for 2, 7, 14, and 28 days, (ii) quantified total, bacterial, and fungal abundances through EL-FAME, PLFA, and qPCR methods, and (iii) measured soil basal respiration. The general dynamic patterns of the three soil microbial fractions in response to soil addition of nutrients and antibiotics were captured by the three methods, which led to strong and positive associations between the abundances of total microorganisms, bacteria, and fungi measured by the three techniques. However, these relationships were found to be stronger between the EL-FAME and PLFA results, indicating that reliability of the fatty-acid based methods is higher than that of the qPCR. Further, soil basal respiration was associated to a higher extent with total, bacterial, and fungal abundances captured by EL-FAME and PLFA analyses than with those measured by qPCR, which suggests that the first two methods are most closely related to the soil living microbial community. In general, dynamics in the abundance of total and bacterial communities were better captured than those of fungi. The PLFA analysis seems to perform better than the EL-FAME method in forest soil and in detecting the small antibiotic-induced decreases in microbial abundances. Since the EL-FAME method is cheaper and allows a much faster processing of samples than the PLFA method, and the reliability of both methods is similar in detecting rapid changes of soil microbial abundances, choosing EL-FAME over PLFA may be advantageous in most cases.

## 1. Introduction

Soil is inhabited by up to 59 % and 90 % of the total known species of bacteria and fungi on Earth, respectively (Anthony et al., 2023). They mediate processes that are key for the many ecosystem services that soil provides: land productivity, nutrient cycling, degradation of contaminants, pathogen control, or climate regulation, among others (Delgado-Baquerizo et al., 2016; Köninger et al., 2022). The quality and quantity of the processes mediated by soil bacteria and fungi are dependent on their diversity and activity, but also on their abundance. In this way, soil multifunctionality has proven to be driven by microbial abundance, among other biotic factors (Wagg et al., 2014; Delgado-Baquerizo et al., 2017). The study of the abundance of bacteria and fungi has been found of interest for soil ecological (Singh and Gupta, 2018; Siles et al., 2023), policy-making (Guerra et al., 2021), agricultural (Wittwer et al., 2021; Hartmann and Six, 2023), and restoration (Plassart et al., 2008; Rodríguez-Berbel et al., 2022) purposes, which demonstrates the multidisciplinary importance of this parameter in soil research.

EL-FAME (ester-linked fatty acid methyl ester), PLFA (phospholipid fatty acid), and quantitative PCR (qPCR) are three of the methods most commonly used for the quantification of bacteria and fungi in soil (Frostegård et al., 1996; Wallander et al., 2013; Mercado-Blanco et al., 2018; Wijaya et al., 2024). EL-FAME and PLFA methods are based on the analysis of the fatty acids of soil microbes, and their further use as representative biomarkers of broad taxonomic groups. The sum of the amounts of every fatty acid assigned to a microbial group provides insights into the soil abundance of that group (Willers et al., 2015). The assignation of fatty acids to the different microbial groups (bacteria, Gram-positive and -negative bacteria, Actinobacteria, fungi, mycorrhiza, etc.) has intensively been investigated and discussed for years (Frostegård et al., 2011; Willers et al., 2015). Nowadays, there is a relative consensus regarding the unspecific microbial biomarkers that can be used to assess total microbial abundance, and the fatty acids that can used as biomarkers of broad taxonomic groups such as bacteria and fungi (McGuire et al., 2013; Willers et al., 2015). The fatty acids used by EL-FAME and PLFA methods as biomarkers coincide, but both approaches methodologically differ, leading to differences in the lipidic fractions that each method captures.

The PLFA method was developed taking advantage of the procedure created by Bligh and Dyer (1959), was firstly applied to soil by Tunlid et al. (1989), and involves the analysis of the fatty acids associated with the phospholipids of the microbial cell membranes. This procedure comprises the (i) extraction of soil total microbial lipids, (ii) fractioning of extracted lipids into neutral lipids, glycolipids, and phospholipids using silicic acid solid-phase extraction and different polarity solvents, (iii) collection and alkaline methanolysis of the phospholipids, and (iv) analysis of the PLFA-ME by gas chromatography (GC). Despite a high-throughput approach has been introduced (Buyer and Sasser, 2012), PLFA analysis is very time-consuming, which is a disadvantage for studies with a large number of samples, especially given the current interest in global- and continental-scale studies (Siles et al., 2023). As an alternative to the PLFA analysis, and by modifying the MIDI (Microbial ID, Inc.) protocol, the EL-FAME method was developed (Ritchie et al., 2000; Schutter and Dick, 2000), which involves: (i) the *in situ* methylation of the microbial ester-linked fatty acids associated to soil microbes, (ii) the partitioning of the resulting EL-FAMEs into an organic phase, and (iii) their further analysis by GC. This method is much less time-consuming, allowing the processing of a high number of samples daily. Furthermore, the EL-FAME method is cheaper (15$ per sample only considering material and reagent costs according to our own calculation) than the PLFA method (30$) and more ecofriendly since less amounts of wastes are generated. Theoretically, while PLFA analysis only captures the fatty acids associated to the phospholipid fraction, the EL-FAME approach addresses fatty acids associated with the three lipid fractions. Since phospholipids have a short turnover time and are consumed rapidly after cell death, the PLFA method is believed to only target the living soil microbial communities, which is the main advantage of the PLFA over the EL-FAME approach (Li et al., 2020).

Different technologies of qPCR (formally called quantitative real-time PCR) exist. One of the most common is the dye-based qPCR, which works with the real-time fluorescence emitted by a dye (e.g., SYBR Green) after it joins the DNA being amplified during the PCR (Rincon-Florez et al., 2013). The fluorescence is detected in real-time by a device, and when the signal overcomes a certain threshold, it is transformed into gene copy numbers on the basis of a standard curve containing the target gene (van Elsas and Boersma, 2011). The tool is compatible with the quantification of microbial groups at different taxonomic levels (domain, phylum, etc.) (Rincon-Florez et al., 2013). Quantification of soil bacteria is conducted by usually targeting the 16S rRNA gene and fungi by analyzing the 18S rRNA gene or the ITS (internal transcribed spacer)1/2 regions (Fierer et al., 2005; Chemidlin Prevost-Boure et al., 2011). The main advantage of the qPCR with respect to the EL-FAME and PLFA analyses lays on its higher specificity; a certain microbial group is more specifically targeted by a pair of primers than by marker fatty acids. However, the relic DNA (Carini et al., 2016; Lennon et al., 2018) and the well-known artifacts associated to PCRs (Qiu et al., 2001) are the main disadvantages of the qPCR analysis.

Studies comparing EL-FAME, PLFA, and qPCR methods are useful to disentangle their reliability for quantifying microbial abundance. Previous studies, such as those of Hinojosa et al. (2005), Miura et al. (2017), Li et al. (2020), and Yu et al. (2021) have compared EL-FAME- and PLFA-based abundances of bacteria and fungi in soils from different ecosystems and under different degrees of anthropogenetic influence, concluding that although both approaches provide consistent results, PLFA data seem to be more sensitive to reflect environmental changes. The EL-FAME and qPCR methods were purposely compared by Pérez-Guzmán et al. (2021), and the PLFA and qPCR approaches by Baldrian et al. (2013), Osburn et al. (2022), and Zhang et al. (2017), among others, demonstrating that, in general, fatty acid-based methods seem to be more reliable than qPCR, although the reliability differences depended on the type of soil and the factor driving the changes in microbial abundance. Despite these works, to the best of our knowledge, there are not studies comparing simultaneous EL-FAME, PLFA, and qPCR methods and considering the temporal dynamics of microbial communities. Most of the aforementioned works considered a sole sampling time.

Soil fungi, and especially bacteria, are characterized by high turnover rates and high responsiveness to environmental changes (Rousk and Bååth, 2007; Randle-Boggis et al., 2018). In this way, when a positive (e.g., soil enrichment with nutrients) or negative (e.g., biocidal agents) stimulus hits a soil ecosystem, bacteria and fungi quickly respond in terms of abundance (Badalucco et al., 1994; Bastida et al., 2013). In short-term scenarios (in the frame of days or weeks), there is a need for methods reliably capturing the response of soil microbial communities to external stimulus in a quantitative manner. Therefore, we aimed here to compare EL-FAME, PLFA, and qPCR methods in an experiment with rapid, sequential changes of microbial multiplication and death, and vice versa. To do that, badland, cropland, and forest soils were amended with nutrients (glucose, ammonium nitrate, and monopotassium) or antibiotics (streptomycin and cycloheximide), and total, bacterial, and fungal abundances were quantified through EL-FAME, PLFA, and qPCR analyses after 2, 7, 14, and 28 days of incubation. We also monitored soil basal respiration throughout the experiment and correlated it with the changes in total, bacterial, and fungal abundances captured by the three approaches in order to get insights into the method better targeting the living fraction of the microbial community. Previous studies have already related soil respiration with the living fraction of soil microorganisms (Blagodatsky et al., 2000; Salazar-Villegas et al., 2016). On one hand, we hypothesized that soil addition of C, N, and P sources would induce an explosive multiplication of total microorganisms, bacteria, and fungi in the range of hours to days after starting the incubation, depending on the soil type. The initially boosted abundances (expected to be captured by the three methods) would gradually decrease at the following sampling times due to the exhaustion of the added nutrients and the competition between microorganisms. On the other hand, we hypothesized that the addition of streptomycin and cycloheximide would have an initially (at 2 or 7 days) detrimental effect on the abundances of soil bacteria and fungi, respectively, leading to the presence of death microbial biomass in soil, with a further increase (at 28 days) in microbial abundances as a consequence of the use of antibiotics as a nutrient source by the surviving microbial community (Badalucco et al., 1994). The reliability of the three methods to detect treatment-driven changes in microbial abundances would decrease in the following order: PLFA > EL-FAME > qPCR, according to their expected capacity to capture only living microorganisms and to not be interfered by dead microbial biomass. EL-FAME and PLFA data would thus correlate stronger between them and with soil basal respiration than qPCR results would do.

## 2. Material and methods

### 2.1. Study sites and soil sampling

Soil from three contrasting ecosystems was collected for the experiment: badland, cropland, and forest. The badland soil was sampled from an area located in the town of Abanilla, Murcia, Spain (38.20459° N, 1.08862° W). The area represents a typical marsh lithology. The climate in the area is semiarid Mediterranean, with an annual average temperature of 19 °C and an annual rainfall of ∼300 mm. At the time of sampling, the area was barely colonized by xerophytic scrubs. The cropland soil was collected from a field located in Albacete, Spain (38.894389° N, 1.988333° W). The field is mainly used for the cultivation of cereals such as maize or wheat, depending on the year and season. The plot is subjected to intensive agriculture with frequent tilling, addition of inorganic fertilizers according to the crop needs, and watering with well water. The climate in the area is typical Mediterranean, with an average annual temperature of ∼15 °C and annual rainfall of ∼350 mm. At the time of sample collection, soil had been recently ploughed, and plants were absent in the parcel. Forest soil was collected from the Regional Park of Sierra Espuña (37.86127° N, 1.54613° W), Murcia, Spain. The vegetation is dominated by pines and holm oaks, among others, with extensive patches of evergreen shrubs. The area is characterized by a semiarid Mediterranean climate, with an annual rainfall of ∼500 mm and an average annual temperature of ∼13 °C at higher elevations (∼1,000 m above sea level), where sampling took place.

For sampling, at each site, an area of approximately 200 m^2^ was delimited, and three equidistant sampling spots were identified and selected to collect three replicate soil samples (one from each spot). Each sample was a composite of five subsamples from the top 20 cm of the soil: four subsamples orthogonally collected in a 2-m radius from a central subsample. Litter, if present, was removed before soil sampling. The three replicate soil samples were then merged into one sample representative of the heterogeneity of each ecosystem type. In total, ∼5 kg of soil were collected from each site. Then, samples were transported to the lab, and subsequently sieved (2 mm mesh), homogenized, and stored at 4 °C until their physicochemical characterization and the setup of the experiment. Soils were physicochemically characterized as described in Siles et al. (2024), and their characterization is presented in Table S1. The three soils presented very contrasting physical, chemical, and biological properties (Table S1).

### 2.2. Experimental setup

The experiment was conducted in 125-mL opaque vials containing 25 g of soil. Microcosms with soil at 60 % of water holding capacity were pre-incubated for 7 days at 25 °C prior to experimental setup. The experiment consisted of three treatments. A “control” treatment that did not receive any external input. A second treatment, designated as “nutrients”, consisted in supplementing soil with C, N, and P in the following chemical forms and concentrations: 2 mg glucose-C g^-1^ soil, 0.1 mg NH_4_NO_3_-N g^-1^ soil, and 0.1 mg KH_2_PO_4_-P g^-1^ soil. These concentrations of nutrients were used in concordance with previous works showing a stimulation of soil microbial abundances after adding these or similar amounts (Demoling et al., 2007; Heuck et al., 2015). A third treatment, designated as “antibiotics”, consisted in adding 25 mg g^-1^ soil of streptomycin sulfate with bactericidal effect (Sigma Aldrich) and 15 mg g^-1^ soil of cycloheximide with fungicidal effect (Thermo Scientific). The used concentrations were higher than those used in other works (Badalucco et al., 1994; Rex et al., 2019) since we were interested in inducing a noticeable detrimental effect on soil bacteria and fungi. Nutrients or antibiotics were added in powder forms to soils and vigorously mixed with a spatula; water content of each soil was then adjusted to 60 % of the water holding capacity with autoclaved distilled water. Afterwards, the microcosms were tightly closed and incubated at 25 °C in the dark. Water loss during incubation was monitored by weighting the microcosms and compensated when required by the addition of sterile water. After 2, 7, 14, and 28 days, the microcosms were destructively sampled, and soil samples were stored at −20 °C until further processing for FAME or PLFA analyses or DNA extraction. In total, the experiment consisted of 144 microcosms: 3 soils (badland, cropland, and forest) × 3 treatments (control, nutrients, and antibiotics) × 4 sampling times (2, 7, 14, and 28 days) × 4 replicates.

### 2.3. Soil basal respiration

For calculation of the soil basal respiration (CO_2_ emissions), the composition of the gas in the microcosms was daily monitored by using a benchtop headspace analyzer CheckMate3 (PBI Dansensor). Percentages of CO_2_ in the atmosphere of the microcosms were transformed into µg C g^-1^ dry weight soil day^-1^ through the ideal gas law according to Hernández and García (2003). The given values of soil basal respiration for each sampling time represent the cumulative respiration during the 4 days prior to sampling expressed on a daily basis; except for the sampling time 2 days, whose soil basal respiration is the cumulative respiration of 2 days.

### 2.4. EL-FAME analysis

Contents of EL-FAME in soil samples were measured according to the method described by Schutter and Dick (2000). Briefly, fatty acids were extracted from microbial cells and released as methyl esters by incubating 3 g of frozen soil with 15 mL 0.2 M methanolic KOH during 1h at 37 °C under periodic shaking. Afterwards, samples underwent pH neutralization with 1 M acetic acid. EL-FAMEs were then partitioned into an organic phase by adding 10 mL hexane and vigorous shaking, followed by centrifugation and evaporation of the hexane in a SpeedVac (Labogene). EL-FAMEs were finally resuspended in isooctane and analyzed with an Agilent 8860 gas chromatograph equipped with a flame ionization detector (FID), using a DB–FastFAME capillary column (30 m × 0.25 mm ID × 0.25 µm film) (Agilent Technologies), with helium as the carrier gas. The chromatographic conditions were as follows: (i) an initial temperature of 80 °C for 1.5 min, (ii) then an increase to 160 °C with a ramp of 40 °C min^-1^, (iii) then to 167 °C at 0.5 °C min^-1^, (iv) then to 200 °C at 30 °C min^-1^, (v) and finally to 230°C at 4 °C min^-1^. Henicosanoic (21:0) methyl ester was used as an internal standard. Fatty acids were identified according to their retention times and their absolute amounts in each sample (nmol g^−1^ dry weight soil) were calculated using commercially available fatty acid methyl ester and bacterial fatty acid methyl ester mixes (Sigma-Aldrich).

Each fatty acid was named according to the following format: total number of C atoms:number of double bonds, followed by additional information about the position of the terminal double bond (ω). Other notations are “Me” for methyl, “cy” for cyclopropane, the prefixes “i” and “a” for iso- and anteiso-branched fatty acids, respectively, and the suffixes “c” and “t” for -cis and -trans configurations, respectively. The fatty acids i15:0, a15:0, i16:0, 16:1ω9c, 10Me16:0, i17:0, cy17:0, 10Me18:0, and cy19:0 were used as bacterial markers. The fatty acids 18:2ω6,9t and 18:2ω6,9c were used as fungal markers. The unspecific microbial fatty acids 14:0, 15:0, 16:0, 17:0, 18:0, and 20:0, along with the bacterial and fungal markers, were used to calculate the total microbial abundance (Joergensen, 2022).

### 2.5. PLFA analysis

Contents of PLFA in soil samples were quantified according to the method developed by Bligh and Dyer (1959). Briefly, microbial lipids were firstly extracted by incubating 3 g of frozen soil with 15 mL of a mixture of chloroform:methanol:citrate buffer (1:2:0.8;v:v:v; citrate buffer, pH 4) during 2h at 20 °C under shaking conditions. After centrifugating (8,000 × g, 10 min), the liquid phase was collected and mixed (by shaking) with chloroform and citrate buffer. The organic phase containing the lipid fraction was collected after centrifugation followed by evaporation of the chloroform in a SpeedVac (Labogene). Lipid classes were then separated by solid phase extraction using the Sep-Pak Silica 3 cc Vac Cartridges (Waters Corp). Each column was firstly conditioned with chloroform. Lipids were then resuspended in chloroform and charged to the columns. Afterwards, column was firstly washed with (i) chloroform, eluting neutral lipids; (ii) secondly with acetone, eluting glycolipids; and (iii) finally with methanol, eluting PLFAs as the fraction of interest. This fraction was collected and then evaporated by using a SpeedVac. PLFA’s methyl transesterification was carried out by incubating PLFAs resuspended in 1 mL of a mixture methanol:toluene (1:1, v:v) with 1 mL of 0.2 M methanolic KOH during 1h at 37 °C. Then, 1 M acetic acid (to stop transesterification reaction), H_2_0 and a mixture hexane:chrolofrom (4:1, v:v) were added, prior to vigorous shaking and centrifugation to collect the hexane phase containing the PLFA-MEs. This action was repeated after the addition a second time of the mixture hexane:chrolofrom. The organic phase was evaporated with a SpeedVac, and the PLFA-MEs were then resuspended in isooctane and analyzed as described for EL-FAME.

### 2.6. Quantitative PCR analysis

DNA from each soil sample was extracted from 250 mg of soil using the DNeasy PowerSoil Pro Kit (QIAGEN) following the manufacturer’s instructions. The DNAs were then spectrophotometrically quality-checked with a NanoDrop (Thermo Fisher Scientific Inc.), fluorometrically quantified with a DS-11 DeNovix device (Life Science Technologies), and their concentrations homogenized to 2 ng µL^-1^ across samples. The abundances of bacterial 16S rRNA and fungal 18S rRNA genes were quantified in DNA extracts using the pairs of primers Eub338/Eub518 for bacteria (Fierer et al., 2005) and FR1/FF390 for fungi (Chemidlin Prevost-Boure et al., 2011). The qPCR reactions were conducted using a QuantStudio 1 system (Applied Biosystems) and SYBR Green as detection system. Each 20μL-reaction contained 10μL PerfeCTa SYBR Green SuperMix Low ROX (2×concentrated, Quantabio), 1 μL each primer (10 μM, Genewiz), 2 μL DNA (2 ng µL^-1^), and 6 μL H_2_O. The thermal conditions were as follows: 95 °C for 3 min followed by 40 cycles of 94 °C for 10 s, 56 °C (bacteria)/58 °C (fungi) for 20 s, and 72 °C for 30 s. In all the cases, after amplification reactions, melting curve and gel electrophoresis analyses were conducted to confirm that the amplified products had the appropriate size. Copy numbers for each gene were calculated using a regression equation for each assay relating the cycle threshold (Ct) value to the known number of gene copies in the standards of a plasmid standard curve containing the appropriate target gene.

The construction of the plasmid standard curves was done as previously reported (Siles and Margesin, 2016). Briefly, a PCR (using the aforementioned primers) copy of the 16S rRNA gene of *Bacillus paralicheniformis* and a copy of the 18S rRNA gene of *Aspergillus fumigatus* were separately cloned using the pGEM®-T Easy Vector System (Promega). Positive clones were selected and subsequently grown overnight, prior plasmid DNA extraction with a QIAprep Spin Miniprep Kit (Qiagen). Plasmid DNAs were quantified and subsequently serially diluted to construct the 16S and 18S rRNA gene standard curves.

Data were expressed as gene copy number g^-1^ dry weight soil and were log-transformed for calculations and graphic representation. QPCR-based total microbial abundances were obtained by summing the 16S and 18S rRNA gene copy numbers. The qPCR efficiencies were 96% (R^2^=0.991) for bacteria and 92% (R^2^=0.989) for fungi.

### 2.7. Statistical analyses

One-way ANOVA (analysis of variance) was applied to determine whether there were significant (P < 0.05) differences between treatments (control, nutrients, and antibiotics) at each sampling time for each soil using the *aov* function of the R package *stats* ver. 4.1.1 (R Core Team). When one-way ANOVA resulted in significant results, Tukey’s HSD (honest significance difference) post-hoc test was used for multiple comparisons of means at a 95% confidence interval. The function *TukeyHSD* and the R packages *multcomp* ver. 1.4-23 (Hothorn et al., 2016) and *multcompView* ver. 0.1-9 (Graves et al., 2015) were used. Normality and heteroscedasticity of data were tested by the Kolmogorov–Smirnov (*ks.test* function, *stats* R package ver. 4.1.1) and Levene tests (*leveneTest* function, *car* R package ver. 3.1-1), respectively. When data did not follow normality and heteroscedasticity assumptions, they were natural log or Box-Cox transformed to make them to satisfy these assumptions. Further, we used linear and quadratic functions to evaluate the strength and shape of the relationships between total, bacterial, and fungal abundances measured by the three methods and between microbial abundances and soil basal respiration. The best model fit was selected by identifying the regression with the lowest Akaike information criterion value. Regressions were calculated using the *stats* R package ver. 4.1.1. Spearman correlation heatmaps were generated using the *Hmisc* ver. 5.1-1 (Harrell Jr, 2019) and *corrplot* ver. 0.92 (Wei et al., 2017) packages in R using untransformed data.

Data visualizations were conducted using the R packages *ggplot2* ver. 3.4.1 (Wickham, 2016) and CorelDRAW ver. 2020.

## 3. Results

### 3.1. Trends of total, bacterial, and fungal abundances measured by EL-FAME, PLFA, and qPCR methods across treatments and soils

EL-FAME, PLFA, and qPCR analyses showed that soil addition of glucose, ammonium nitrate, and monopotassium phosphate greatly increased total, bacterial, and fungal abundances in badland, cropland, and forest soils (Fig. 1). Except for the badland soil, a relatively good consistency between the percentages of increment measured by the three methods was found (Table S2). This consistency was higher for the results obtained by the EL-FAME and PLFA methods, which captured averagely increasing rates of 100 % across soils and times for the three microbial fractions. Amounts of total, bacterial, and fungal PLFAs represented ∼27, 38, and 15 % of the EL-FAMEs, respectively, across all the samples in our experiment (Table S3). These percentages for the three microbial fractions were always higher in forest soil than in the other two soils (Table S3). The EL-FAME and PLFA methods coincided in showing the highest levels of total, bacterial, and fungal abundances in badland and cropland soil after 2 days of incubation with decreasing abundances afterwards (Fig.1). This trend was also detected by qPCR for total and bacterial communities but not for fungi. In the forest soil, the highest amounts of total microorganisms, bacteria, and fungi were determined by EL-FAME and PLFA methods after 7 days of incubation, while qPCR detected them after 14 days (Fig. 1). A chronological decoupling between fatty acid and DNA approaches was thus observed. At 28 days, the final incubation time, the three methods detected significantly increased amounts of total microorganisms and bacteria in the nutrient-amended treatment with respect to the control. In the badland soil, EL-FAME and qPCR-based fungal abundances were higher in the nutrient addition treatment, but this was not captured by PLFA. The three methods agreed in showing not significant differences in quantities of fungi between control and nutrient treatment in the cropland soil (Fig. 1). The PLFA and qPCR methods did not detect increased abundances of bacteria and fungi in forest soil at 28 days in the nutrient addition treatment with respect to the control, while EL-FAME did so for fungi.

**Fig. 1.**
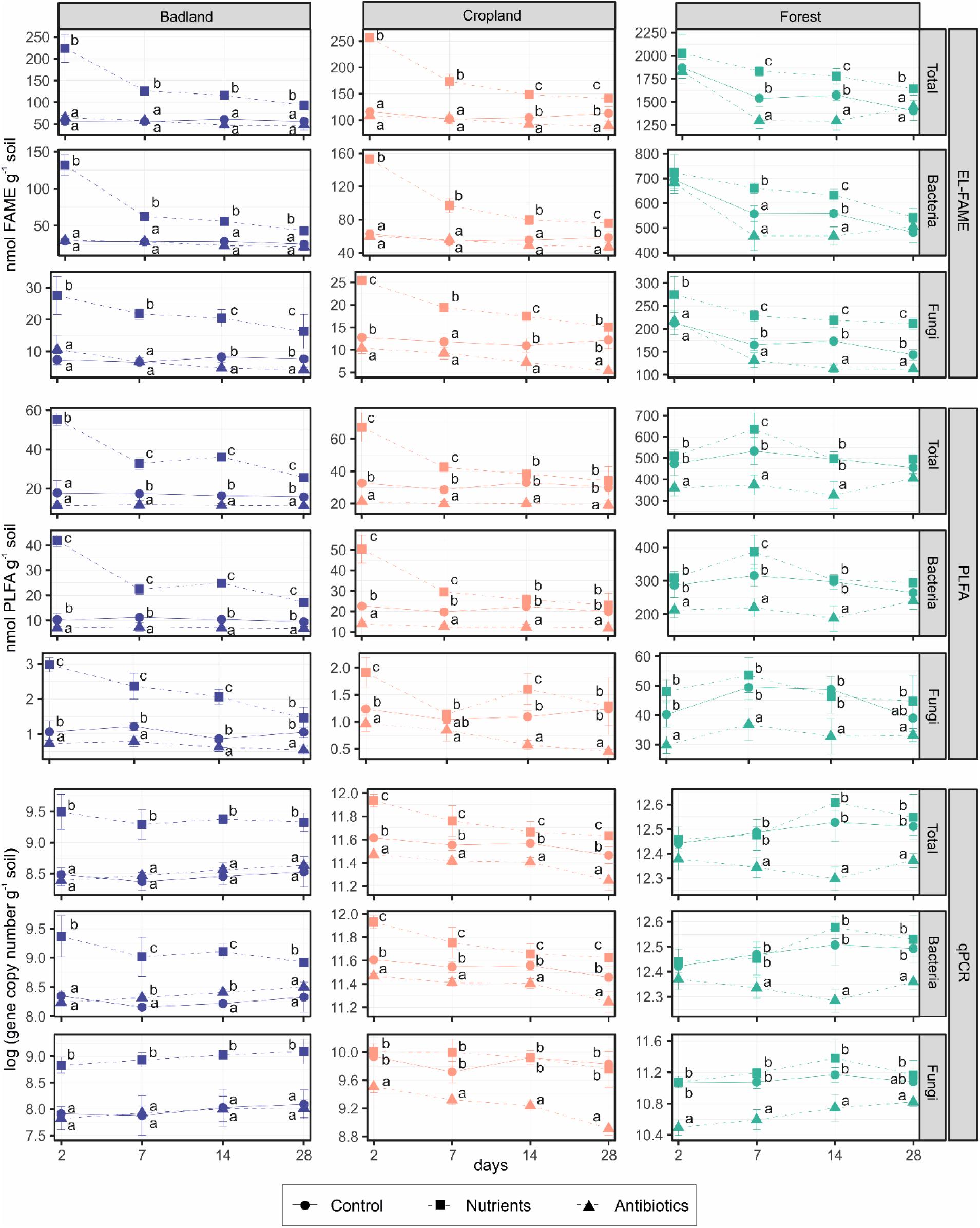
Total, bacterial, and fungal abundances measured by EL-FAME, PLFA, and qPCR methods in badland, cropland, and forest soils at 2, 7, 14, and 28 days after not addition (control) or addition of nutrients or antibiotics. Bars represent standard desviations. For each soil, microbial fraction, method, and sampling time, different letters indicate significance of differences at P < 0.05 according to ANOVA and Tukey’s HSD test.

In quantitative terms, the impact of soil addition of streptomycin sulfate and cycloheximide was smaller than that provoked by nutrient addition (Fig. 1 and Table S2). The average decreasing percentages of total, bacterial, and fungal abundances were 17, 18, and 31% according to the EL-FAME analysis, respectively; 32, 34, and 36% according to the PLFA method; and 32, 13, and 69 % according to qPCR across the three soils and times analyzed. In general, the detrimental effect was the strongest on fungi, in badland soil, and after 14 days of incubation (Fig. 1 and Table S2). Antibiotic addition decreased amounts of total microorganisms, bacteria, and fungi in the three soils according to the PLFA method at all the times; however, this trend was not that clearly detected by EL-FAME and qPCR methods, especially in the badland soil (Fig. 1). The three methods were more sensitive in detecting the antibiotic-driven changes in microbial abundances in cropland and forest soils than in badland soil. Increased microbial abundances at the final incubation time were not measured for any microbial fraction.

### 3.2. Relationships between the total, bacterial, and fungal abundances measured by EL-FAME, PLFA, and qPCR methods

Regression (Fig. 2) and correlation (Fig. 3) analyses considering all the samples in the study demonstrated a relatively good consistency between the results obtained with the three methods. When the three methods were pairwise compared for each microbial fraction, total, bacteria, and fungal abundances were significantly positively associated. The relationships between EL-FAME- and PLFA-based abundances of total microorganisms, bacteria, and fungi were stronger than those found between EL-FAME and qPCR and between PLFA and qPCR (Fig. 2, Fig. 3). EL-FAME and qPCR results for the three microbial fractions were related to a higher extent than PLFA and qPCR results. The relationships between methods for the three microbial fractions were stronger in badland and cropland soils than in forest soil (Fig. S1, Fig. S2).

**Fig. 2.**
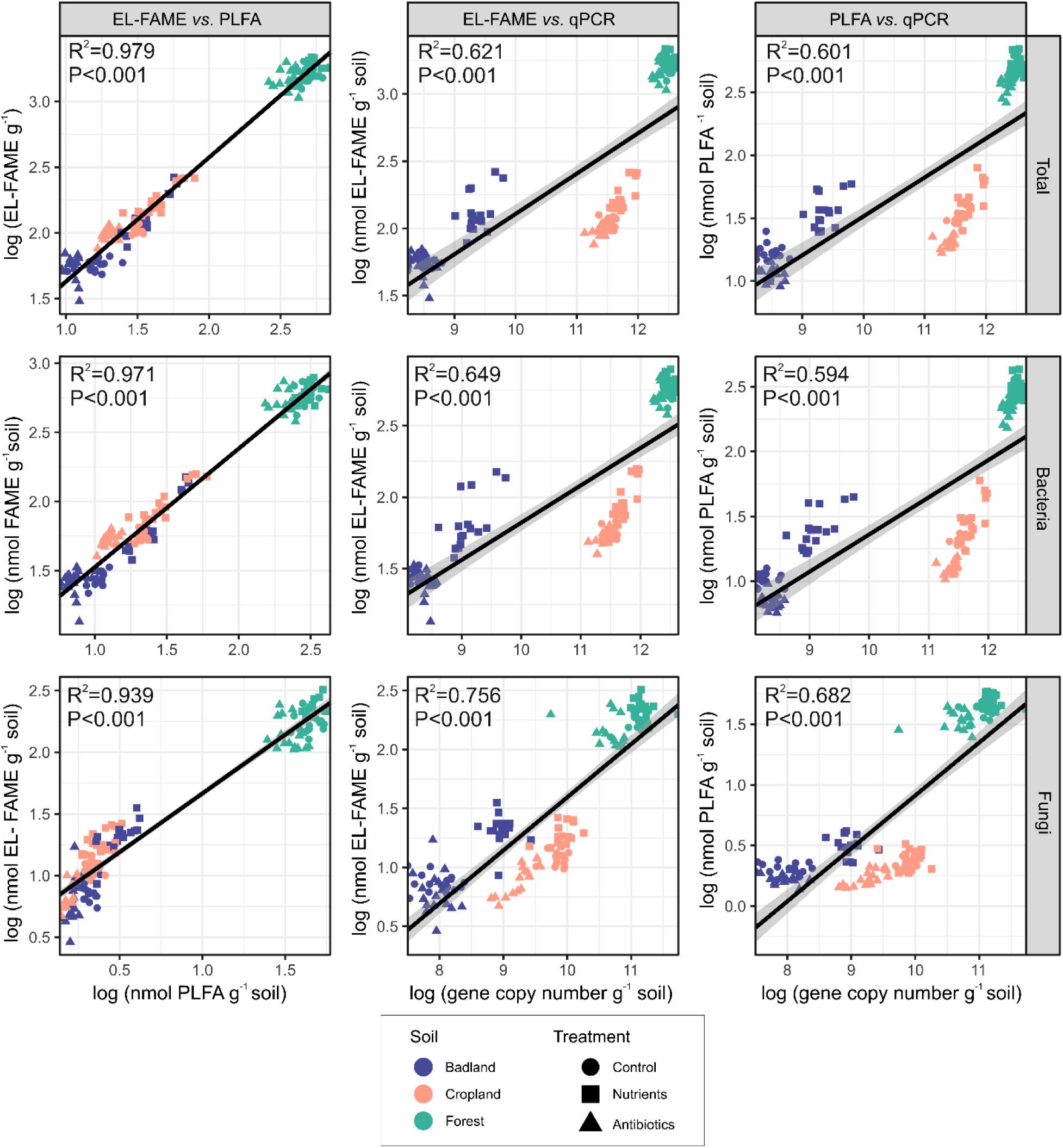
Regression analyses relating total, bacterial, and fungal abundances measured by EL-FAME, PLFA, and qPCR methods in badland, cropland, and forest soils at 2, 7, 14, and 28 days after not addition (control) or addition of nutrients or antibiotics (n=144). Shaded areas represent 95 % confidence intervals for the regression line. R^2^ and p-values are shown for each regression analysis. Data are log-transformed. All the regressions fitted with the linear model.

**Fig. 3.**
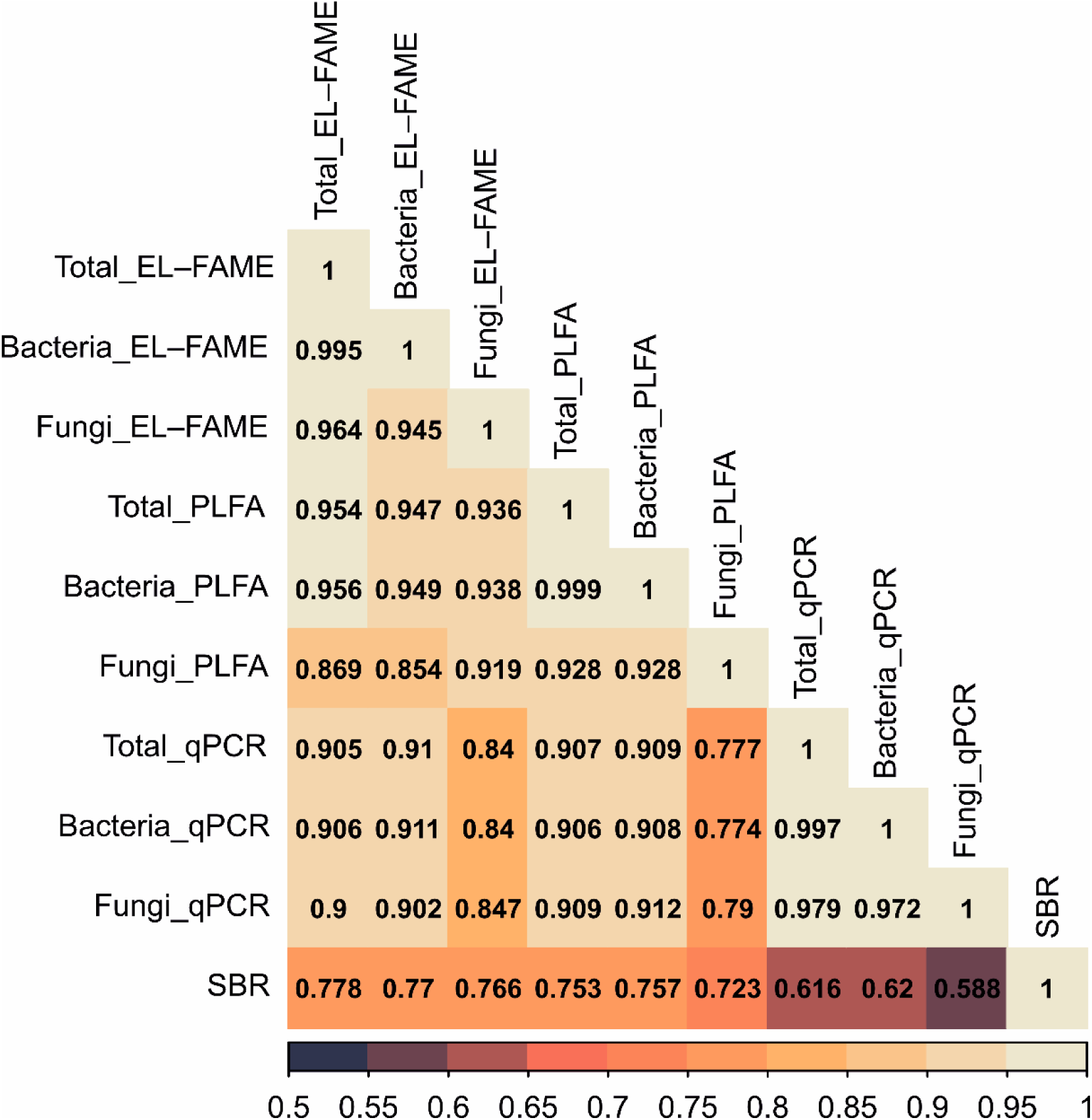
Heatmap showing significant (P < 0.05) Spearman correlations between total, bacterial, and fungal abundances measured by EL-FAME, PLFA, and qPCR methods and between them and soil basal respiration in badland, cropland, and forest soils at 2, 7, 14, and 28 days after not addition (control) or addition of nutrients or antibiotics (n=144). Non-transformed data were used for Spearman correlations.

### 3.3. Soil basal respiration and its relationships with the total, bacterial, and fungal abundances measured by EL-FAME, PLFA, and qPCR methods

Soil nutrient addition stimulated soil basal respiration by ∼50, 131, and 3 times with respect to the control treatment in the badland, cropland, and forest soils, respectively, after 2 days. (Fig. S3 and Table S2). These increments were proportionally decreasing over time until the final sampling time at 28 days, when soil respiration in the nutrient addition treatment was 130 and 85% higher than in the control in the badland and cropland soil, respectively, or did not significantly differ with respect to the control in the forest soil (Fig. S3). The effect of the antibiotic addition on soil basal respiration was soil dependent. In badland and forest soils, addition of antibiotics decreased soil basal respiration after 7 (∼47 and 19 % decreasing in badland and forest soil, respectively) and 14 (∼70 and 18 % decreasing in badland and forest soil, respectively) days with respect to the control, but increased (by ∼30 and 50 % in badland and forest soil, respectively) it after 28 days (Fig. S3 and Table S2). In cropland soil, the antibiotic addition significantly increased soil basal respiration at all the sampling times, with the highest increment rates (of ∼337 %) being detected 7 days after antibiotic addition (Fig. S3 and Table S2).

When the three soils were altogether considered, soil basal respiration was significantly positively associated with total, bacterial, and fungal abundances (Fig. 3 and Fig. 4). The relationships found between soil basal respiration and FAME- and PLFA-based abundances for the three microbial fractions were much stronger than those found between soil respiration and qPCR-based abundances (Fig. 3 and Fig. 4). The relationships between soil basal respiration and FAME- and PLFA-based abundances for the three microbial fractions followed a unimodal pattern. Instead, an inverse unimodal curve was the best describing the relationships between soil respiration and amounts of total microorganisms, bacteria, and fungi determined by qPCR (Fig. 3 and Fig. 4). The strength of the associations between the total microbial abundance measured with the three methods and soil respiration did not greatly differ from that obtained for the relationships between bacterial and fungal abundances determined by the three methods and soil respiration. When the associations between the abundances of the three microbial fractions and soil respiration were separately analyzed for each soil, EL-FAME- and PLFA-based total, bacterial, and fungal abundances of the badland and forest soils were significantly related to soil basal respiration, which was not found in the cropland soil (Fig. S3). The qPCR results for the three microbial fractions were found to correlate with the soil basal respiration in the badland soil, but not in cropland and forest soils.

**Fig. 4.**
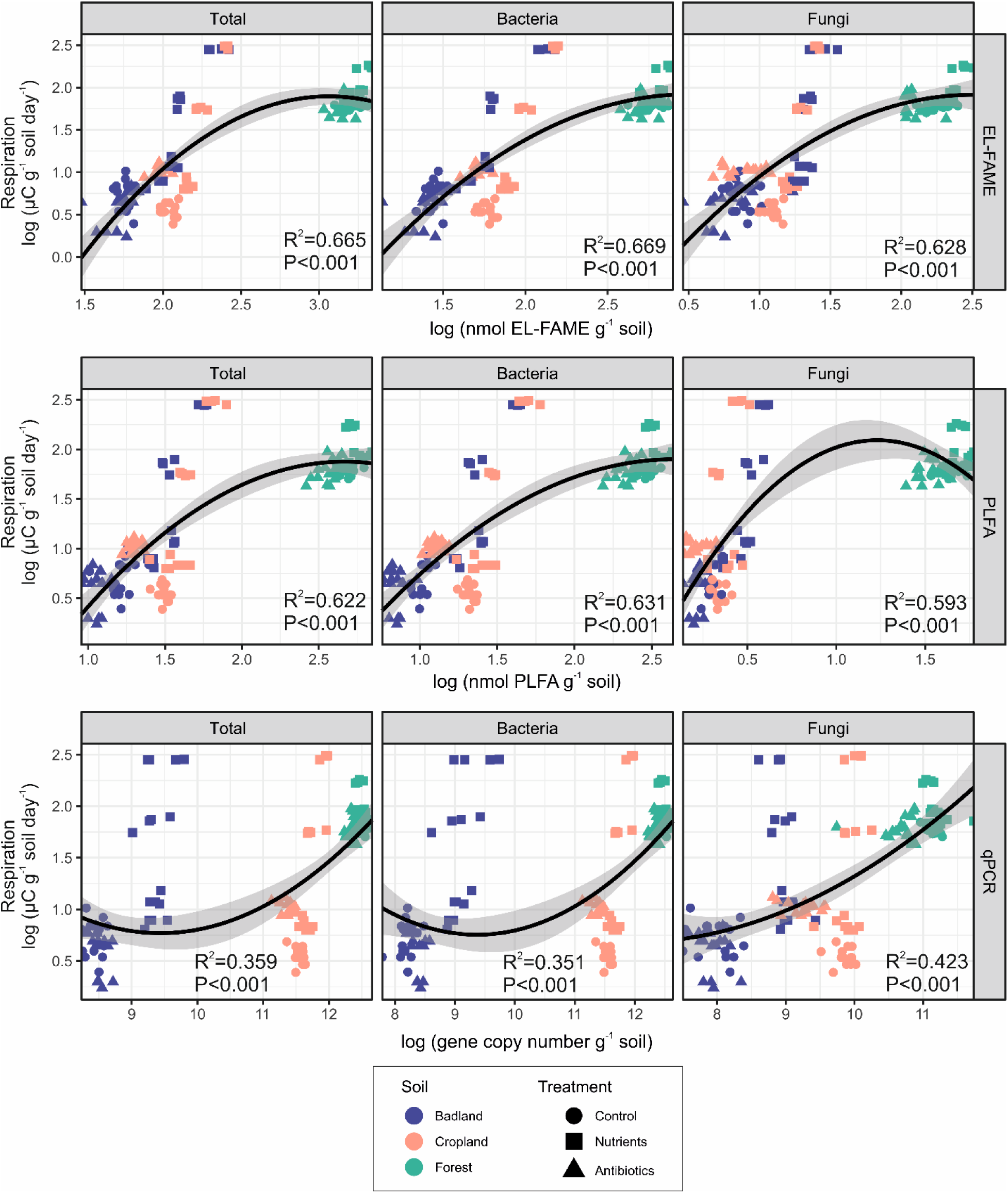
Regression analyses relating soil basal respiration and total, bacterial, and fungal abundances measured by EL-FAME, PLFA, and qPCR methods in badland, cropland, and forest soils at 2, 7, 14, and 28 days after not addition (control) or addition of nutrients or antibiotics (n=144). Shaded areas represent 95 % confidence intervals for the regression line. R^2^ and p-values (P) are shown for each regression analysis. All the regressions fitted with the unimodal model.

## 4. Discussion

The present study represents a step forward in the field of soil microbial ecology by (i) comparing simultaneously the reliability of EL-FAME, PLFA, and qPCR methods, (ii) using a short-term experiment with quick changes in microbial abundances in response to soil addition of nutrients and biocidal compounds; and (iii) relating changing multi-approach-based abundances of total microorganisms, bacteria, and fungi with soil basal respiration. The three methods were able to capture the broad patterns in the soil microbial dynamics produced by the addition of nutrients and antibiotics. Despite this, fatty acid-based methods showed higher degree of reliability than qPCR. It has always been accepted that selecting the EL-FAME method over the PLFA-based one involves a loss in the reliability of the measured microbial abundances, it is the price to pay in order to speed up the number of samples processed per day. However, our results demonstrate that the sensitivity of both methods to detect rapid changes in the amounts of soil total microorganisms, bacteria and fungi is similar. This information is of interest for researchers designing a short-term experiment and looking for the most suitable cost-benefit approach for the measurement of soil microbial communities or for scientists considering stablishing a new analytical method in their labs.

### 4.1. Relationships between total, bacterial, and fungal abundances measured by EL-FAME, PLFA, and qPCR methods

The ideal approach for researchers to assess soil microbial abundances should be fast, cost-effective, reliable, able to only target the living or active fraction of the community, and sensitive to microbial turnover at short-term, independently of the soil type. With their pros and cons, EL-FAME, PLFA, and qPCR are nowadays three of the most common methods used in soil science to monitor the abundance of soil microbial abundance (Wijaya et al., 2024). As an example, a search on Scopus on April 2024 with the terms “soil” and “EL-FAME/FAME”, “PLFA”, or “qPCR” gives 536, 2,657, and 2,762 results, respectively. Attempts to transform data of abundance provided by EL-FAME, PLFA, or qPCR analyses into absolute amounts of soil microbial biomass (i.e., µg microbial biomass C g^-1^ soil) have been made (Baldrian et al., 2013; Willers et al., 2015). However, the reliability of these conversion factors has been questioned (Willers et al., 2015) and further research is needed for verification (Camenzind et al., 2024). Therefore, the three methods used here quantify soil microbial abundance and are a proxy of soil microbial biomass.

Overall, we found that the abundances of total microorganisms, bacteria, and fungi determined by the three methods were strongly associated, contrary to our initial expectations. EL-FAME, PLFA, and qPCR-based abundances of total microorganisms and bacteria correlated stronger than fungal ones. This is seen in relation to the higher number of fatty acid biomarkers used for the assessment of total and bacterial abundances, which facilitates capturing the changes happening in the entire microbial community. Further, the associations between the different multi-approach-based microbial abundances were stronger in badland and cropland soils than in forest soil. In the forest soil, the EL-FAME results for the three microbial fractions were associated with the qPCR and PLFA data weaker than the results of these last two did. This is probably a consequence of using the highly-alkaline (pH∼13) KOH methanolic solution for the in-situ methylation of the fatty acids in the EL-FAME method in the forest soil, with a high content in soil organic matter and plant material. This could favour the dispersal of soil colloids and increase the extraction of fatty acids from plant material and humified soil organic matter, interfering with the quantification of real amounts of bacteria and fungi (Li et al., 2020). The lower resolution that the EL-FAME method has in the quantification of bacteria and fungi in forest soils with respect to the PLFA approach has previously been reported (Yu et al., 2021). Despite this, it is important to highlight that, in our study, the EL-FAME data significantly correlated with the PLFA and qPCR data and that the EL-FAME method was able to successfully capture the impact that both nutrients and antibiotics produced on soil microbial abundances.

PLFA was anticipated to be the method best capturing the expected microbial dynamics since microbial cell membrane phospholipids are believed to quickly degrade once cell dies through the action of phospholipases and phosphodiesterases, among others (Zhang et al., 2019). However, the much better resolution of the PLFA method to capture the microbial dynamics with respect to FAMES and qPCR, demonstrated by a low association of data from the different methods, was not clearly observed. In fact, the microbial dynamics captured by PLFA and EL-FAME after nutrient and antibiotic addition were quite similar, and even similar percentages of microbial growth were obtained. This would indicate that the turnover of the phospholipids analyzed through the PLFA analysis did not greatly differ from that of the total lipids (phospholipids, glycolipids, and neutral lipids) analyzed by the EL-FAME method. Zhang et al. (2019) estimated that turnover of phospholipids occurs in a period ranging from 14 to 27 h after amending a paddy soil with free microbial phospholipids. However, it has been argued that turnover of phospholipids in real conditions may be slower (Frostegård et al., 2011) as phospholipids are not free in soil, and the process of phospholipid degradation is highly dependent on environmental conditions like pH and soil texture (Vachon et al., 2021; Joergensen, 2022). We considered here three soils with very different physicochemical properties and textures, and similar patterns were found in the three soils. Unfortunately, degradation times of neutral lipids (usually found in microbial storage structures, especially in fungi, and in recently-formed bacterial necromass), and of glycolipids (membrane lipids whose functions are not yet fully understood but recently related to conditions of environmental stress) in soil are not known, to the best our knowledge, and should be object of study in the future (Lekberg et al., 2022; Gorka et al., 2023). Despite this, our results do not indicate that neutral lipids and glycolipids remain for long periods of time (more than 5 days) in soil upon cell death. It may be argued that the good correlation we found between EL-FAME and PLFA data is because phospholipids were the dominant lipid fraction in our EL-FAME analysis; however, across all the samples in our experiment, total, bacterial, and fungal PLFAs only represented ∼27, 38, and 15 % of the EL-FAMEs, respectively. These values are a bit lower than those found in other studies (Miura et al., 2017; Li et al., 2020; Yu et al., 2021), although the variability between studies is high, which is explained by interlaboratory differences in protocols and analytical conditions.

Despite the good agreement between the three methods for the three microbial fractions, we noticed some differences that are worth noting. For instance, microbial abundances did not follow the pattern we hypothesized after soil addition of biocides. In general, amounts of total microorganisms, bacteria and fungi decreased immediately after addition of antibiotics in the three soils, but none of them increased after 14 or 28 days of incubation despite other works have reported this phenomenon (Badalucco et al., 1994). This is probably a consequence of the relatively high concentrations of streptomycin (inhibits the initiation of protein synthesis in the prokaryotic community and causes misreading of messenger RNA) and cycloheximide (inhibits the peptidyl transferase activity of the eukaryotic 60S ribosomal subunit) used (Badalucco et al., 1994), which makes that longer period of times are probably needed to detect an increase in microbial abundances (Chen et al., 2023). We also noticed that the PLFA method was more sensitive than EL-FAME and qPCR analyses to capture the discrete detrimental effect of antibiotics on bacteria and fungi abundance after two days of incubation, which postulates PLFA as the method to choose for the monitoring of the microbial abundances in very short-term experiments after application of a discrete negative stimulus to a soil.

The qPCR data across the experiment correlated relatively well with the data of the fatty-acid-based methods, but these associations were weaker than those found between the EL-FAME- and PLFA-based abundances for total microorganisms, bacteria, and fungi. This result was expected and is explained by the quicker degradation of fatty acids than DNA in soil. Carini et al. (2016) quantified in 40% the amount of “relic DNA” averagely presents in a soil sample and that can thus interfere with the quantification of the actual microbial abundance, especially in short-term experiments with rapid, sequential changes in microbial abundances. We observed that the highest amounts of total microorganisms, bacteria, and fungi were captured by the EL-FAME and PLFA methods after 7 days of incubation in the forest soil after nutrient addition, while the qPCR detected them after 14 days. This may be indicating that nutrients induced initially (after 7 days of incubation) in this soil an increase of the microbial cell size (or in storage structures) that resulted in a rise in soil fatty acid contents without a concomitant increase in copies of ribosomal genes. In a subsequent stage (after 7 days), the nutrients would have induced a multiplication of the microbial communities, which implies DNA replication and an increase in the number of copies of ribosomal genes. In relation to this, it has been reported that cell fatty acid content is proportional to its size (Ehlers et al., 2010), and that bacteria can reduce their cell size by three times in response to nutrient starvation (Vadia et al., 2017). Furthermore, we noticed that the C:N:P ratio in the forest soil (368:15:1) was much more unbalanced than that in the badland (8:0.8:1) and cropland (19:1.4:1) soils, which could explain why this phenomenon happened in forest soil and not in the badland and cropland soils since Ehlers et al. (2010) already reported a short-time enlargement of microbial cells after nutrient addition in a highly P-limited soil.

### 4.2. Soil basal respiration and its relationship with microbial abundances measured by EL-FAME, PLFA, and qPCR methods

We here monitored the soil basal respiration in order to know how the activity of soil microbial communities changed with the addition of nutrients and antibiotics and get insights on how treatment-mediated shifts in microbial respiration and abundances relate. We expected that respiration and microbial abundance would change in parallel since both parameters have been demonstrated to usually correlate (Holden and Treseder, 2013; Chen et al., 2019; Siles et al., 2022). Especially since we here measured soil basal respiration (cumulative respiration expressed on a daily basis) and not soil respiration at a given moment (Demoling et al., 2007; Creamer et al., 2014). The expected associations were expected to give us insights into the ability of each method to only capture the living fraction of the microbial community and their capacity to not be interfered by dead microbial biomass (Blagodatsky et al., 2000). Our regression and correlation analyses proved that total, bacterial, and fungal abundances and soil basal respiration were significantly and positively related across our experiment. Overall, the strength of the associations of the EL-FAME- and PLFA data with soil basal respiration were similar for the three microbial fractions, and higher than those found between the qPCR data and soil basal respiration. This demonstrates, as above discussed, that using the EL-FAME method does not imply an important loss in sensitivity to measure the real amounts of total microorganisms, bacteria, and fungi in an experiment with rapid changes in microbial abundances. This assumption would apply for the three microbial fractions since similar association were found between soil basal respiration and the three microbial groups. The weaker correlations of soil basal respiration with the qPCR results for the three microbial fractions are explained by the soil extraction of relic DNA (usually small fragments bonded to inorganic and organic substances and highly present in determined soil microhabitats) that interferes in the quantification of the living microbial community (Carini et al., 2016; Lennon et al., 2018).

Further, total, bacterial, and fungal abundances determined by the three methods (not for qPCR results in the forest soil) positively correlated with soil basal respiration in badland and forest soils, but not in cropland soil. In this soil, no significant correlations were found between microbial abundances and soil basal respiration. We explain this as a result of the increased basal respiration detected in the cropland soil after the addition of the antibiotics, which occurred without a parallel increase of microbial abundance. In fact, microbial abundances and soil basal respiration were significantly positively correlated when analyses were re-done without including the data for the microcosms with antibiotics (Fig. S4). Previous studies have argued that the lack of decline or increase in soil basal respiration after soil addition of biocides may have three different explanations: (i) the added biocidal compounds are used as substrate due to the presence of, for example, antibiotic-resistance genes; (ii) the quick biocidal effects on the target group release nutrients previously locked in biomass that are used for the survival microbes; or (iii) biocides alleviate biotic competition and increase resource availability for the unaffected microorganisms (Rousk et al., 2009; Shawver et al., 2024). Under these circumstances, soil microorganisms seem to prioritize activity over multiplication.

## 5. Conclusions

The three methods, EL-FAME, PLFA, and qPCR, were able to detect the broad pattern of changes in total, bacterial, and fungal abundances induced by the soil addition of nutrients and antibiotics in the conducted short-term experiment. However, the stronger associations found between the PLFA and EL-FAME results across the experiment and between these and the soil basal respiration demonstrate that these two methods are more sensitive than qPCR in capturing changes in the abundance of soil microbial communities. Choosing fatty acid-based methods over qPCR is advised in experiments where a high sensitivity in quantifying the living soil bacteria and fungi is sought. The PLFA method seems to perform better than the EL-FAME analysis in the forest soil and in detecting the very short-term, reduced antibiotic-induced decreases in microbial abundances. Since the EL-FAME method is much faster, allowing the analysis of a high number of samples daily, cheaper, and eco-friendly (less wastes are generated) in comparison with the PLFA approach, and the reliability of both methods is similar, choosing EL-FAME over PLFA is recommended in most of scenarios.

## CRediT authorship contribution statement

**José A. Siles:** Conceptualization, Formal analysis, Investigation, Visualization, Writing - Original Draft. **Roberto Gómez-Pérez:** Investigation, Writing - review & editing. **Alfonso Vera:** Investigation, Writing - review & editing. **Carlos García**: Funding acquisition, Resources, Writing - review & editing. **Felipe Bastida**: Conceptualization, Funding acquisition, Resources, Writing - review & editing.

## Acknowledgements

This publication is part of the I+D+I project PID2020-114942RB-I00 funded by MCIN/AEI/10.13039/501100011033. This publication is part of the I+D+I project CPP2022-009734 funded by MCIN/AEI/10.13039/501100011033 and the European Union-NextGenerationEU/PRTR. This study formed part of the AGROALNEXT programme and was supported by MCIN with funding from European Union NextGeneration EU (PRTR-C17.I1) and by Fundación Séneca with funding from Comunidad Autónoma Región de Murcia (CARM). J.A.S. acknowledges the support of the grant IJC2018-034997-I funded by MICIU/AEI/10.13039/501100011033.

