## Supplementary material for "A comparison among EL-FAME, PLFA, and quantitative PCR methods to detect changes in the abundance of soil bacteria and fungi"

for

The supplementary material includes Tables S1-S3 and Fig. S1-S4.

**Table S1.** Main physicochemical properties of the three soils used in the present study. Values are means (n=3) ± standard deviations. Different letters within each row indicate significance of differences at $P < 0.05$ according to ANOVA and Tukey's HSD test. EC, electrical conductivity; SOM, soil organic matter; C, carbon; N, nitrogen; P, K, S, Ca, and Mg, phosphorus, potassium, sulfur, calcium, and magnesium, respectively.

| | Badland | Cropland | Forest |
|---|---|---|---|
| Clay (%) | 57.5 | 31.6 | 3.7 |
| Silt (%) | 36.1 | 32.1 | 15.9 |
| Sand (%) | 6.4 | 36.3 | 80.4 |
| Texture | Clay | Clay Loam | Loamy Sand |
| pH | 8.72±0.02 b | 8.93±0.01 c | 8.29±0.03 a |
| EC (μS $cm^{-1}$) | 486.0±31.4 | 176.3±7.8 a | 309.7±20.0 |
| Total C (g $kg^{-1)}$ | 73.13±1.03 b | 66.36±0.04 a | 172.9±8.8 c |
| Organic C (g $kg^{-1}$) | 5.70±0.78 a | 13.51±3.65 b | 172.9±4.8 c |
| SOM (%) | 0.98±0.13 a | 2.33±0.63 b | 29.88±0.84 c |
| Total N (g $kg^{-1}$) | 0.55±0.04 a | 1.01±0.02 b | 7.25±0.33 c |
| C/N | 10.38±1.68 a | 13.39±1.40 b | 23.86±0.49 c |
| Total P (g $kg^{-1}$) | 0.70±0.04 b | 0.70±0.06 b | 0.47±0.06 a |
| Total K (g $kg^{-1}$) | 12.10±0.61 b | 11.13±0.01 b | 8.80±0.46 a |
| Total S (g $kg^{-1}$) | 1.90±0.10 b | 1.43±0.06 a | 1.70±0.10 b |
| Total Ca (g $kg^{-1}$) | 169.3±9.54 c | 128.0±3.46 b | 98.4±6.56 a |
| Total Mg (g $kg^{-1}$) | 12.67±0.21 a | 11.77±0.45 a | 16.03±0.81 b |
| Total microbial abundance (nmol FAME g-1 soil) | 56.78±7.6 a | 115.83±8.3 b | 1869.3±114.9 c |
| Bacterial abundance (nmol FAME $g^{-1}$ soil) | 29.14±3.5 a | 62.94±5.1 b | 693.7±33.1 c |
| Fungal abundance (nmol FAME $g^{-1}$ soil) | 7.40±1.9 a | 12.78±0.4 b | 213.4±25.6 c |

**Table S2.** Percentages of increasing or decreasing (negative values) EL-FAME, PLFA, and qPCR-based total, bacterial and fungal abundances induced by nutrient or antibiotic addition with respect to the control in badland, cropland, and forest soils at 2, 7, 14, and 28 days after not addition (control) or addition of nutrients or antibiotics. Only increasing or decreasing percentages for treatments showing significant differences with respect to the control are shown.

| | | | | EL-FAME | | | PLFA | | | qPCR | | |
|---|---|---|---|---|---|---|---|---|---|---|---|---|
| Soil | Treatment | Time | Respiration | Total | Bacteria | Fungi | Total | Bacteria | Fungi | Total | Bacteria | Fungi |
| Badland | Nutrients | 2 | 4869 | 295.2 | 352.6 | 272.1 | 211.4 | 309.6 | 180.8 | 1061.4 | 1186.3 | 857.7 |
| Badland | Antibiotics | 2 | | | | | | -32.10 | -30.90 | | | |
| Cropland | Nutrients | 2 | 13005 | 121.8 | 143.1 | 99.18 | 105.4 | 123.3 | 54.61 | 110.4 | 112.4 | 20.00 |
| Cropland | Antibiotics | 2 | 310.5 | | | -18.87 | -34.66 | -38.30 | -21.87 | -28.28 | -27.54 | -62.48 |
| Forest | Nutrients | 2 | 193.1 | | | 28.89 | | | | | | |
| Forest | Antibiotics | 2 | | | | | -24.09 | -25.84 | -25.91 | | | -63.01 |
| Badland | Nutrients | 7 | 887.7 | 124.6 | 120.7 | 227.8 | 89.05 | 101.1 | 95.00 | 789.1 | 796.5 | 778.4 |
| Badland | Antibiotics | 7 | -47.37 | | | | -33.40 | -33.81 | -35.36 | | 45.34 | |
| Cropland | Nutrients | 7 | 2315 | 69.73 | 80.76 | 64.73 | 47.94 | 49.84 | | 67.56 | 67.13 | |
| Cropland | Antibiotics | 7 | 336.9 | | | | -30.77 | -36.37 | | -27.22 | -26.70 | -61.18 |
| Forest | Nutrients | 7 | 30.76 | 18.90 | 18.70 | 38.54 | 19.19 | 22.53 | | | | |
| Forest | Antibiotics | 7 | -19.66 | -15.68 | | -20.27 | -29.78 | -30.34 | -25.62 | -26.94 | -25.33 | -66.22 |
| Badland | Nutrients | 14 | 276.0 | 90.46 | 93.87 | 148.3 | 120.7 | 140.5 | 139.3 | 714.9 | 698.6 | 736.0 |
| Badland | Antibiotics | 14 | -70.00 | | | | -30.79 | -32.06 | -27.67 | | 55.24 | |
| Cropland | Nutrients | 14 | 160.0 | 42.10 | 43.63 | 59.46 | | | 46.52 | 27.11 | 27.73 | |
| Cropland | Antibiotics | 14 | 245.0 | -12.32 | | -34.23 | -39.12 | -44.43 | -47.41 | -31.09 | -29.97 | -79.25 |
| Forest | Nutrients | 14 | 23.48 | 13.12 | 13.40 | 26.36 | | | | | | |
| Forest | Antibiotics | 14 | -17.83 | -17.71 | -16.31 | -34.75 | -34.36 | -36.84 | -32.98 | -41.52 | -40.63 | -60.86 |
| Badland | Nutrients | 28 | 130.4 | 64.54 | 71.88 | 113.1 | 63.32 | 83.60 | | 482.5 | 248.8 | 884.9 |
| Badland | Antibiotics | 28 | 30.43 | | | | -29.63 | -27.74 | -48.58 | | | |
| Cropland | Nutrients | 28 | 84.62 | 25.16 | 30.11 | | | | | 46.71 | 48.01 | |
| Cropland | Antibiotics | 28 | 230.8 | -20.86 | -19.25 | -56.46 | -36.01 | -40.09 | -64.03 | -39.11 | -37.94 | -87.96 |
| Forest | Nutrients | 28 | 27.4 | 16.96 | | 47.48 | | | | | | |
| Forest | Antibiotics | 28 | 49.1 | | | -21.50 | | | | -27.46 | -26.46 | |

**Table S3.** Proportion (average±standard deviation) of measured amounts of PLFAs with respect to FAMEs for total microorganisms, bacteria, and fungi.

| Soil type | Treatment | Time (days) | Total (%) | Bacteria (%) | Fungi (%) |
|---|---|---|---|---|---|
| Badland | Control | 2 | 30.99±9.00 | 34.82±6.65 | 14.35±3.06 |
| Badland | Antibiotics | 2 | 17.29±2.50 | 23.21±1.50 | 7.78±2.78 |
| Badland | Nutrients | 2 | 24.95±2.43 | 31.92±2.30 | 11.10±1.92 |
| Cropland | Control | 2 | 28.32±1.76 | 36.08±2.90 | 9.69±1.01 |
| Cropland | Antibiotics | 2 | 19.77±2.53 | 23.36±3.00 | 9.40±1.83 |
| Cropland | Nutrients | 2 | 26.10±2.99 | 32.99±4.45 | 7.49±0.85 |
| Forest | Control | 2 | 25.42±3.96 | 41.51±6.57 | 19.05±3.20 |
| Forest | Antibiotics | 2 | 19.70±2.36 | 31.31±3.70 | 13.78±1.52 |
| Forest | Nutrients | 2 | 25.20±1.54 | 42.79±2.23 | 17.61±1.26 |
| Badland | Control | 7 | 31.20±4.34 | 39.81±5.05 | 18.62±4.01 |
| Badland | Antibiotics | 7 | 19.98±3.40 | 25.98±5.73 | 11.76±2.77 |
| Badland | Nutrients | 7 | 26.00±2.34 | 36.01±4.21 | 10.87±1.47 |
| Cropland | Control | 7 | 28.13±2.23 | 36.88±2.87 | 8.91±0.86 |
| Cropland | Antibiotics | 7 | 19.31±1.42 | 22.57±1.44 | 9.09±1.45 |
| Cropland | Nutrients | 7 | 24.57±2.86 | 30.66±2.85 | 5.91±0.69 |
| Forest | Control | 7 | 34.70±4.43 | 56.80±5.67 | 30.01±2.81 |
| Forest | Antibiotics | 7 | 29.42±7.14 | 48.24±12.20 | 28.51±7.23 |
| Forest | Nutrients | 7 | 34.80±4.94 | 58.77±9.09 | 23.52±3.38 |
| Badland | Control | 14 | 27.09±2.35 | 36.03±3.54 | 10.79±2.52 |
| Badland | Antibiotics | 14 | 24.67±6.37 | 31.51±7.18 | 13.01±2.85 |
| Badland | Nutrients | 14 | 31.26±1.99 | 44.52±3.43 | 10.14±0.76 |
| Cropland | Control | 14 | 31.82±3.25 | 40.54±3.70 | 10.14±1.80 |
| Cropland | Antibiotics | 14 | 22.04±2.98 | 25.43±3.30 | 8.40±2.44 |
| Cropland | Nutrients | 14 | 25.77±1.17 | 32.27±1.83 | 9.18±1.60 |
| Forest | Control | 14 | 31.62±3.11 | 53.44±5.14 | 28.19±2.84 |
| Forest | Antibiotics | 14 | 25.47±6.47 | 40.78±10.47 | 29.25±8.93 |
| Forest | Nutrients | 14 | 27.93±1.80 | 48.01±2.71 | 21.26±3.03 |
| Badland | Control | 28 | 27.82±3.12 | 37.70±5.06 | 13.94±3.35 |
| Badland | Antibiotics | 28 | 25.11±10.83 | 34.28±15.80 | 13.37±5.08 |
| Badland | Nutrients | 28 | 27.95±4.47 | 40.37±5.16 | 9.92±4.21 |
| Cropland | Control | 28 | 26.78±2.52 | 34.50±3.08 | 10.60±4.16 |
| Cropland | Antibiotics | 28 | 21.51±2.22 | 25.43±2.39 | 8.38±0.54 |
| Cropland | Nutrients | 28 | 24.22±6.16 | 30.62±7.64 | 8.51±3.37 |
| Forest | Control | 28 | 32.52±2.94 | 55.29±5.01 | 27.36±4.76 |
| Forest | Antibiotics | 28 | 28.03±3.26 | 47.65±5.37 | 29.60±3.59 |
| Forest | Nutrients | 28 | 30.28±5.36 | 54.69±10.10 | 21.09±3.65 |
| | | Overall | 26.60±4.36 | 37.97±9.96 | 15.02±7.53 |

**Fig. S1.** Regression analyses relating total, bacterial, and fungal abundances measured by EL-FAME, PLFA, and qPCR methods in badland, cropland, and forest soils at 2, 7, 14, and 28 days after not addition (control) or addition of nutrients or antibiotics. Shaded areas represent 95 % confidence intervals for the regression line. $R^2$ and p-values (P) are shown for each regression analysis. Data are log transformed. All the regressions fitted with the linear model.

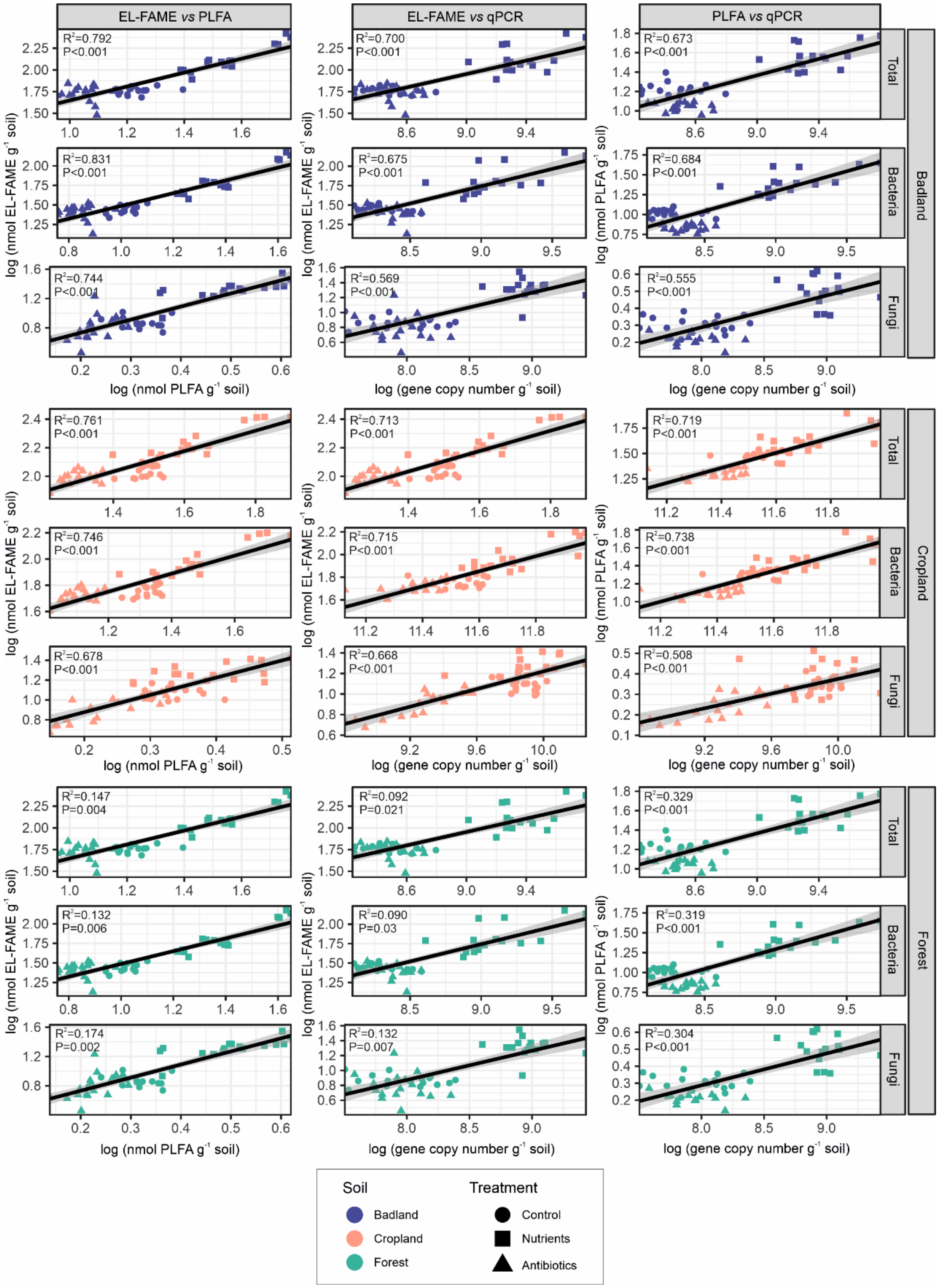

**Fig. S2.** Heatmap showing significant (P < 0.05) Spearman correlations between total, bacterial, and fungal abundances measured by EL-FAME, PLFA, and qPCR methods and between them and soil basal respiration (SBR) in (a) badland, (b) cropland, and (c) forest soils at 2, 7, 14, and 28 days after not addition (control) or addition of nutrients or antibiotics.

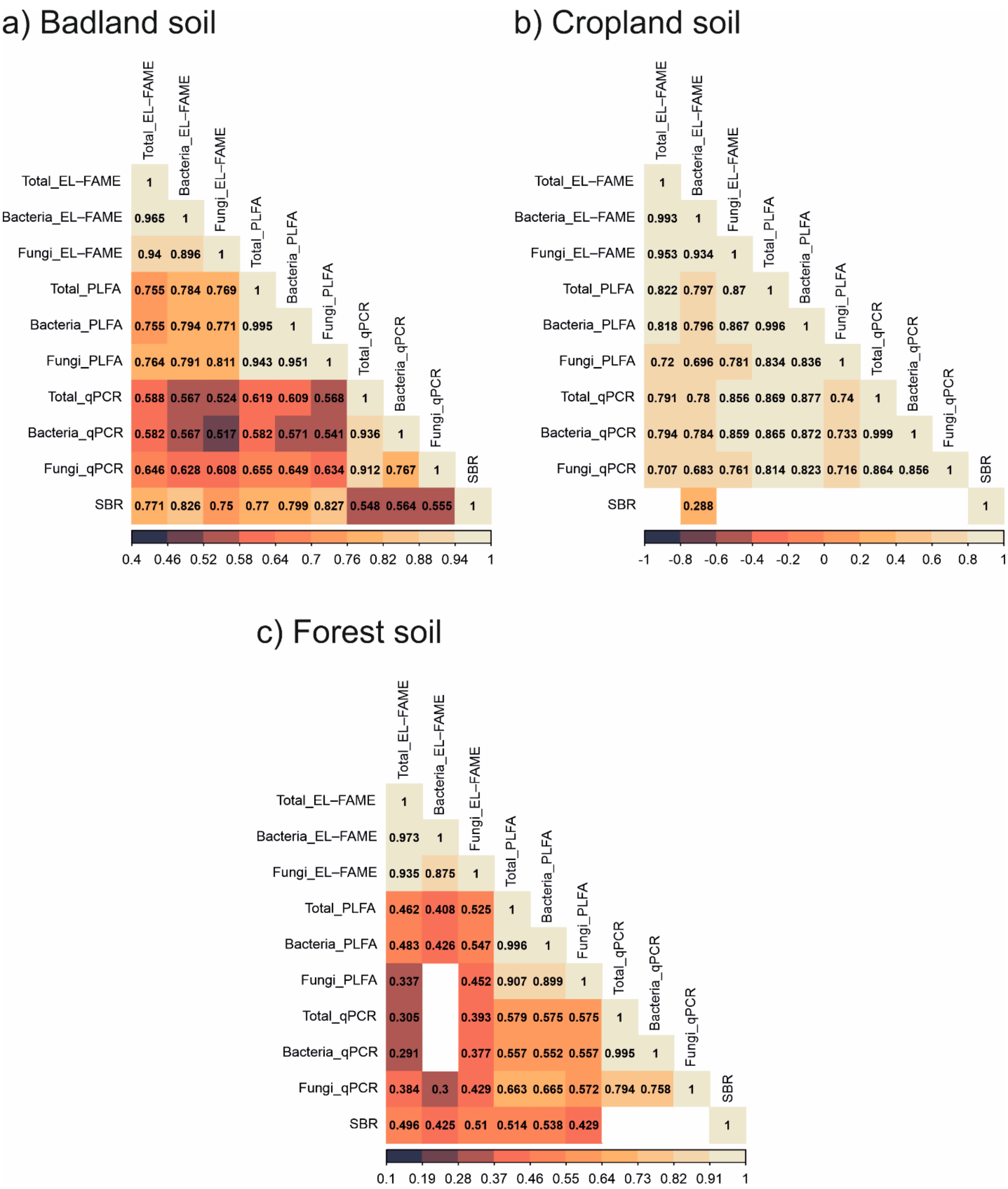

**Fig. S3.** Soil basal respiration in badland, cropland and forest soils measured at 2, 7, 14, and 28 days after not addition (control) or addition of nutrients or antibiotics. Bars represent standard deviation. For each soil and sampling time, different letters indicate significance of differences at $P < 0.05$ according to ANOVA and Tukey's HSD test.

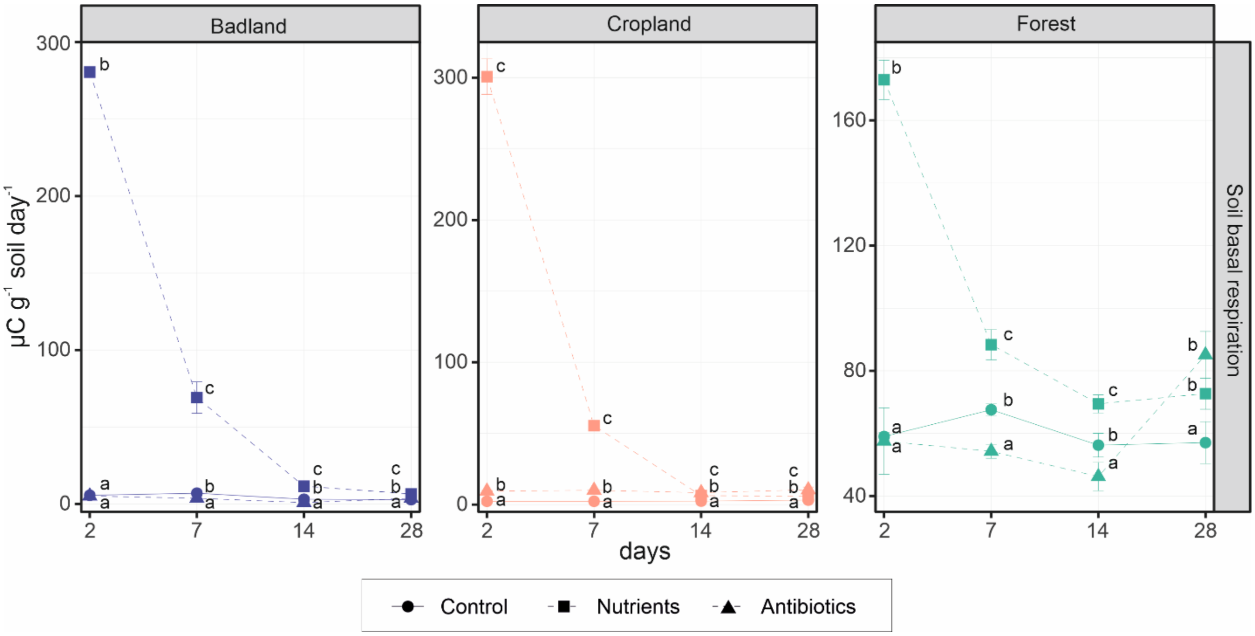

**Fig. S4.** Heatmap showing significant ($P < 0.05$) Spearman correlations between soil basal respiration (SBR) and total, bacterial, and fungal abundances measured by EL-FAME, PLFA, and qPCR methods in the cropland soil at 2, 7, 14, and 28 after not addition (control) or addition of nutrients.

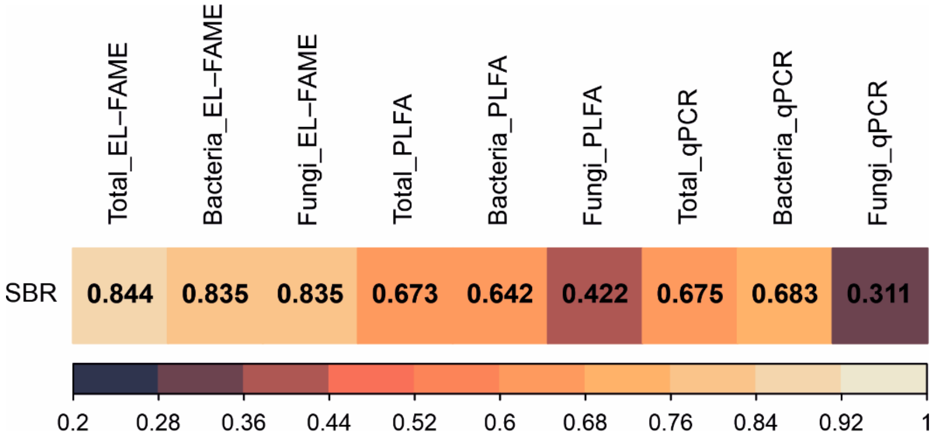
